# Template-assisted synthesis of adenine-mutagenized cDNA by a retroelement protein complex

**DOI:** 10.1101/344556

**Authors:** Sumit Handa, Yong Jiang, Sijia Tao, Robert Foreman, Raymond F. Schinazi, Jeff F. Miller, Partho Ghosh

**Affiliations:** Department of Chemistry & Biochemistry, University of California, San Diego, CA 92093, USA; Center for AIDS Research, Laboratory of Biochemical Pharmacology, Department of Pediatrics, Emory University School of Medicine, Atlanta, GA 30322, USA; Departments of Microbiology, Immunology, and Molecular Genetics, Molecular Biology Institute, and California NanoSystems Institute, University of California, Los Angeles, Los Angeles, CA 90095, USA

## Abstract

Diversity-generating retroelements (DGRs) create unparalleled levels of protein sequence variation through mutagenic retrohoming. Sequence information is transferred from an invariant template region (*TR*), through an RNA intermediate, to a protein-coding variable region. Selective infidelity at adenines during transfer is a hallmark of DGRs from disparate bacteria, archaea, and microbial viruses. We recapitulated selective infidelity *in vitro* for the prototypical *Bordetella* bacteriophage DGR. A complex of the DGR reverse transcriptase bRT and pentameric accessory variability determinant (Avd) protein along with DGR RNA were necessary and sufficient for synthesis of template-primed, covalently linked RNA-cDNA molecules, as observed *in vivo*. We identified RNA-cDNA molecules to be branched and most plausibly linked through 2′-5′ phosphodiester bonds. Adenine-mutagenesis was intrinsic to the bRT-Avd complex, which displayed unprecedented promiscuity while reverse transcribing adenines of either DGR or non-DGR RNA templates. In contrast, bRT-Avd processivity was strictly dependent on the template, occurring only for the DGR RNA. This restriction was mainly due to a noncoding segment downstream of *TR*, which specifically bound Avd and created a privileged site for processive polymerization. Restriction to DGR RNA may protect the host genome from damage. These results define the early steps in a novel pathway for massive sequence diversification.

## INTRODUCTION

Diversity-generating retroelements (DGRs) create massive protein sequence variation in ecologically diverse bacteria, archaea, and microbial viruses, including constituents of the human microbiome (1–7). An extraordinary prevalence of DGRs was recently identified in the microbial ‘dark matter’ (8). Variation by DGRs extends to at least 10^20^ possible sequences (9). The only parallel to this scale of variation occurs in vertebrate adaptive immune systems, in which, for example, 10^14-16^ sequences are possible in antibodies and T cell receptors (10,11). Strikingly, DGRs dedicate 10^3^-fold less DNA but can achieve 10^4-6^-fold greater variation than these adaptive immune systems. Massive protein sequence variation enables binding of novel targets, and is recognized to enable the immune system to adapt to dynamic environments.

A similar fitness benefit appears to be provided by the prototypical DGR of *Bordetella* bacteriophage (1). This DGR encodes the phage’s receptor-binding protein Mtd (12,13), which is localized to the tip of phage tail fibers (Fig. S1A). Massive sequence variation in Mtd (~10^13^) enables the phage to adapt to the loss of *Bordetella* surface receptors due to environmentally-responsive shifts in gene expression or immune-driven selection (4,14). For example, the *Bordetella* adhesin pertactin, which is expressed by the *in vivo* pathogenic or Bvg^+^ phase of *Bordetella*, can serve as a surface receptor for the phage (1,15). However, when *Bordetella* transitions to the *ex vivo* or Bvg^−^ phase, expression of pertactin is lost. The loss of this phage receptor is counterbalanced by the DGR through the generation of Mtd variants, some of which have the ability to maintain phage infectivity by binding surface receptors expressed by Bvg^−^ *Bordetella* (1,15).

DGRs diversify proteins through mutagenic retrohoming, which requires three fundamental components: a specialized reverse transcriptase and two nearly identical repeat regions, a variable region (*VR*) and a template region (*TR*) (Fig. 1A). *VR* is located within the coding sequence of a diversified protein, such as *mtd*, and *TR* is proximally located in a noncoding region. Sequence information is transferred unidirectionally from *TR* to *VR* in a selectively error-prone manner focused on adenines in *TR*. Adenine-specific mutagenesis is a unique hallmark of DGRs. The process involves transcription of *TR* to produce *TR*-RNA, which is then reverse transcribed by the DGR reverse transcriptase to produce *TR*-cDNA (1,2,5,16–19) (Fig. 1A). The *TR*-cDNA, which has specific errors due to the unfaithful copying of adenines, homes to and replaces *VR* through a mechanism that is as yet undefined. As a result of the adenine-specificity of mutagenesis, only amino acids that are encoded with adenines in *TR* are subject to diversification. The most frequently observed adenine-containing codon in *TR* is AAY, which encodes Asn. Adenine-mutagenesis of AAY can result in the encoding of 14 other amino acids, which cover the gamut of amino acid chemical character, but cannot result in a stop codon (12). Notably, adenine-encoded variable amino acids have been seen to form the solvent-exposed, ligand-binding sites of diversified proteins and participate in binding ligands (9,12,15,20).

**Figure 1.**
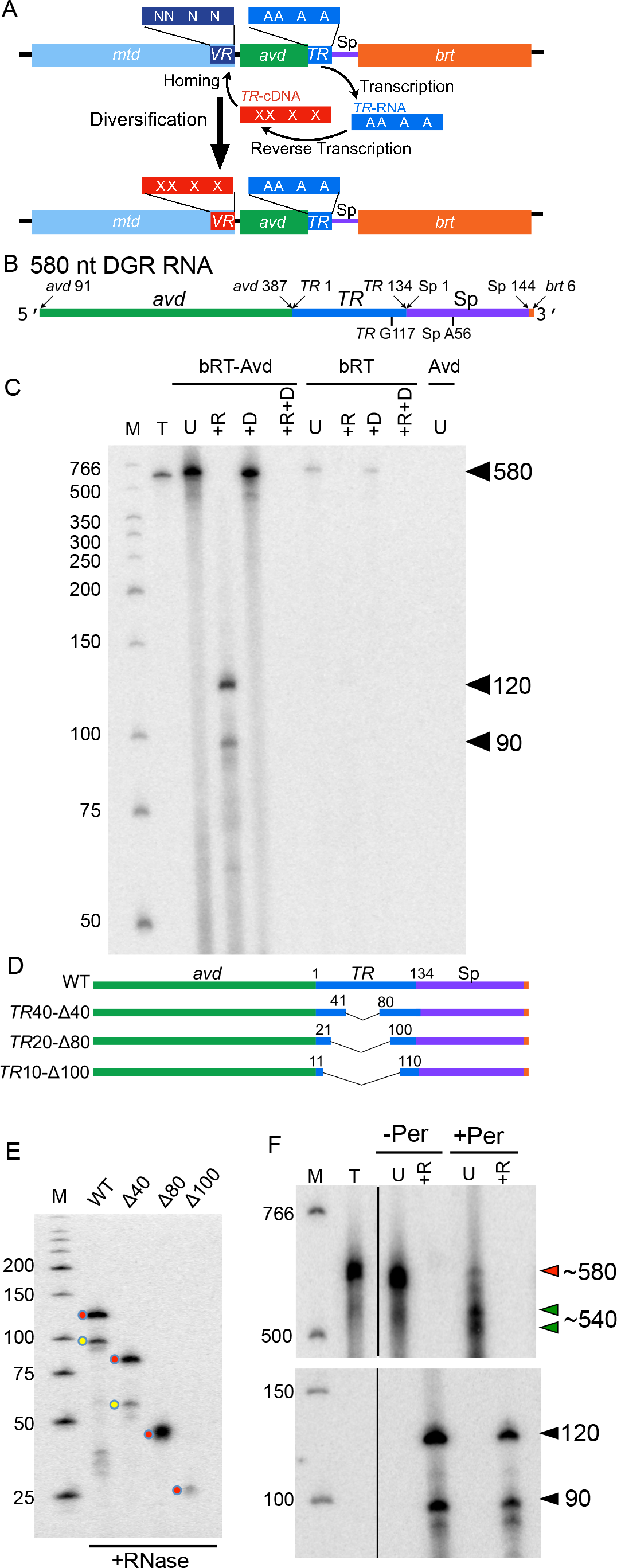
*In vitro* template-primed cDNA synthesis. (A) *Bordetella* bacteriophage DGR diversification of Mtd. *mtd* contains a variable region (*VR*), which encodes the receptor-binding site of the Mtd protein. Downstream of *VR* is the template region (*TR*). Adenines in *TR* (“A”) are frequently replaced by another base in *VR* (“N”). *TR* is transcribed to produce *TR*-RNA, which is then reverse transcribed to *TR*-cDNA. During this process, adenines in *TR* are mutagenized, as depicted by “X” in *TR*-cDNA. Adenine-mutagenized *TR*-cDNA homes to and replaces *VR*, resulting in diversification of Mtd. *bRT* is the DGR reverse transcriptase, and *avd* the DGR accessory variability determinant. (B) Sequence elements of the 580 nt DGR RNA template used for reverse transcription reactions. (C) bRT-Avd, bRT, or Avd was incubated with the 580 nt DGR RNA and dNTPs, including [α-^32^P]dCTP, for 2h. Products resulting from the incubation were untreated (U), or treated with RNase (+R), DNase (+D), or both RNase and DNase (+R+D), and resolved by 8% denaturing polyacrylamide gel electrophoresis (PAGE). Lane T corresponds to internally-labeled 580 nt DGR RNA as a marker for the size of the template. The positions of the 580 nt band, and 120 and 90 nt cDNA bands are indicated. Nuclease-treated samples were loaded at twice the amount as untreated samples, here and throughout unless otherwise indicated. Lane M here and throughout corresponds to radiolabeled, single-stranded DNA molecular mass markers (nt units). (D) DGR RNA templates containing internal truncations in *TR*. (E) Radiolabeled cDNA products resulting from bRT-Avd activity for 2 h with intact (WT) or internally truncated 580 nt DGR RNA as template. Samples were treated with RNase and resolved by denaturing PAGE. The positions of the 120 and 90 nt cDNAs produced from intact template are indicated by red and yellow circles, respectively, as are positions of the correspondingly shorter cDNAs produced from truncated RNA templates. (F) Radiolabeled products resulting from bRT-Avd activity for 2 h with the 580 nt DGR RNA as template. Prior to reverse transcription, the RNA template was mock-treated (−Per) or treated with periodate (+Per). Products of the reaction were untreated (U) or treated with RNase (+R), and resolved by 4% (top) or 8% (bottom) denaturing PAGE. In the top gel, the red arrowhead indicates the ~580 nt species, and the green arrowheads the several ~540 nt species. In the bottom gel, the black arrowheads indicate the 120 and 90 nt cDNA products. The black vertical line within the gel indicates irrelevant lanes that were removed for display purposes. A two-fold higher quantity was loaded for +Per samples than −Per samples.

In a number of retrotransposons, reverse transcription is primed by the DNA target of integration, thereby coupling cDNA synthesis to homing (21,22). However, recent *in vivo* studies show that cDNA synthesis by the *Bordetella* bacteriophage DGR does not require the target *VR*, and is instead primed by the DGR RNA template itself (19). Reverse transcription to produce *TR*-cDNA requires the *Bordetella* bacteriophage DGR reverse transcriptase bRT and the DGR accessory variability determinant (Avd) protein (1,19,23). The structure of bRT is not known, but it is predicted to resemble the reverse transcriptase domain of group II intron maturases (24–27). bRT appears to contain no additional functional domains. The structure of Avd is known, and it forms a pentameric and highly positively charged barrel (23). Avd interacts functionally with bRT (23).

To gain mechanistic insight into cDNA synthesis and adenine-mutagenesis by DGRs, we reconstituted reverse transcription by the *Bordetella* bacteriophage DGR *in vitro*. We found that a purified complex of bRT and Avd along with the DGR RNA template were necessary and sufficient for the synthesis of template-primed and covalently linked RNA-cDNA molecules, as observed *in vivo* (19). We show that these RNA-cDNA molecules were branched due to a priming site located internally in the RNA, and linked most plausibly through 2’-5’ RNA-cDNA phosphodiester bonds. bRT alone had no reverse transcriptase activity, and required association with Avd for this function. Most significantly, we found that adenine-mutagenesis was intrinsic to the bRT-Avd complex, which displayed an unprecedented promiscuity in reverse transcribing adenines of either DGR or non-DGR RNA templates. The processivity of bRT-Avd was strictly dependent on the RNA template, occurring only for the DGR RNA. This restriction was mainly due to a noncoding region downstream of *TR*, which specifically bound Avd and created a privileged site for processive polymerization by bRT-Avd. These results provide fundamental advances in understanding the mechanism of mutagenic retrohoming.

## MATERIAL AND METHODS

### Cloning, Expression, and Purification of Proteins

*Bordetella* bacteriophage bRT, with an N-terminal His_6_-tag followed by a PreScission protease cleavage site, and Avd were co-expressed using pET-28b (Novagen) and pACYCDuet-1 vectors (Novagen), respectively, in *Escherichia coli* BL21-Gold (DE3). Bacteria were grown with shaking at 37 °C to OD_600_ 0.6−0.8, and induced with 0.5 mM isopropyl β−D-1-thiogalactopyranoside (IPTG), and then grown further overnight with shaking at 20 °C. Bacteria were centrifuged and frozen at −80 °C, and then thawed and re-suspended in 500 mM (NH_4_)_2_SO_4_, 50 mM HEPES, pH 8.0, 0.1% β-mercaptoethanol, 20% glycerol, 2 mM PMSF, protease inhibitor tablet(s) (Roche), DNase I (Ambion, 0.25 units/mL) and RNase A/T1 mix (Ambion, 0.2 μL/mL). Bacteria were lysed using an Emulsiflex C-5, and the lysate was clarified by centrifugation. His_6_-bRT/Avd from the supernatant was purified by Ni^2+^-NTA column chromatography. The His_6_-tag was cleaved with PreScission protease, and the bRT-Avd complex was further purified by gel filtration chromatography (Superdex 200). Two peaks were evident, with the first peak corresponding to the bRT-Avd complex and the second to bRT alone. Avd was expressed and purified as described previously (*19*).

### RNA Synthesis and Purification

RNA was synthesized *in vitro* through runoff transcription reactions performed overnight at 37 °C using 40 ng/μL DNA template, 4 mM NTPs (Promega), 0.5 units/μL RNase inhibitor (NEB), and 0.6 mg/mL T7 RNA polymerase in T7 buffer (3 mM MgCl_2_, 5 mM DTT, 2 mM spermidine, 40 mM Tris, pH 7.5). Products were extracted with phenol:chloroform:isoamyl alcohol (25:24:1 v/v, P:C:I) and ammonium acetate/ethanol (AA/E) precipitated. The pellet was resuspended in water and digested with 0.5 units/μL DNase I (Ambion) for 1 h at 37 °C, and then P:C:I extracted and AA/E precipitated as above. RNA was further purified by denaturing polyacrylamide gel electrophoresis. RNA was recovered from the gel by diffusion overnight in 300 mM NaCl, 0.1 % SDS, 1 mM EDTA, and extracted with P:C:I and chloroform:isoamyl alcohol (24:1 v/v, C:I), and AA/E precipitated. The pellet was washed with cold 80% ethanol, air dried for 10 min at RT, and resuspended in water. Internally-labeled RNA was produced similarly, except the NTP was 0.5 mM and 0.2 μCi/μL of [α-^32^P]UTP was included. Mutations were introduced into DNA templates by the QuikChange method (Agilent). Scrambled sequences in the Sp region are indicated in Table S1. For the non-DGR RNA template, pET28a was digested with HindIII and used for *in vitro* runoff transcription from its T7 promoter.

### Reverse Transcription

Reactions were carried out in 20 μL containing 1.8 μM bRT-Avd, bRT, or Avd, 100 ng/μL RNA template, 100 μM dNTPs (Promega) (or varying concentrations of certain dNTPs), 0.5 μCi/μL [α-^32^P]dCTP, 20 units RNase inhibitor (NEB) in 75 mM KCl, 3 mM MgCl_2_, 10 mM DTT, 50 mM HEPES, pH 7.5, 10 % glycerol for 2 h at 37 °C. Oligodeoxynucleotide-primed reactions included 0.25-1 μM primer (Table S2). Samples were digested with proteinase K (Ambion, 0.4 mg/mL) at 50 °C for 30 min, and quenched with 2 mM EDTA. Samples were then desalted using a G-25 spin column, ammonium acetate/ethanol precipitated with 10 μg glycogen included with the ethanol (AA/E-glycogen). The pellet was resuspended in 10 μL water. Nuclease treatments were carried out on reverse transcription products (2 μL for RNase digestion and 1 μL for DNase digestion) in 10 μL containing 0.5 μL RNase A/T1 mix or 0.5 μL DNase I in RNase or DNase buffer. For digestion with UDG and RNase, 10 μL containing 2 μL of reverse transcription products, 5 units UDG (NEB), and 2 μL RNase A/T1 in UDG reaction buffer was incubated at 37 °C for 20 min, and quenched by the addition of 10 μL denaturing loading dye.

Samples were heated for 5 min at 95 °C in denaturing loading dye, resolved on denaturing polyacrylamide gels, and visualized by autoradiography using a Typhoon Trio (GE Healthcare Life Sciences). Quantification of band intensities was carried out using ImageQuant TL 8.1 (GE Healthcare Life Sciences). Background, as measured from a lane containing no sample in the gel being quantified, was subtracted from band intensities.

Reverse transcription reactions with HIV-1 RT were carried out as above, except in 10 μL and containing 10 units RNase inhibitor (NEB), 0.1 μCi/μL [α-^32^P]dCTP, 30 ng/μL RNA template, 1 μM P^G117^ primer (Table S2) and 2 units of HIV-1 RT (Worthington Biochemical), and the reaction was carried out for 30 min.

### Periodate Treatment of RNA

RNA (200 μg) was incubated in 400 μL borax buffer, pH 9.5 (50 mM boric acid and 5 mM borax) and 50 mM sodium periodate at 30 °C for 1 h in the dark, followed by addition of 40 μL glycerol, and quenching of the reaction at RT for 10 min. The sample was concentrated using a Speedvac to 50 μL, then diluted to 400 μL in borax buffer, pH 9.5, and incubated at 45 °C for 90 min. The sample was finally purified through a desalting column (G-25) and a denaturing polyacrylamide gel, as above.

### RT-PCR of RNA-cDNA molecules

RNA-cDNA molecules (200 ng) resulting from bRT-Avd activity on DGR RNA was annealed with in 13 μL with primer 1 (Table S2) at 65 °C for 5 min in the presence of 0.5 mM dNTPs and 5% DMSO. The sample was quenched on ice for 5 min and 200 units Super RT (BioBharati Life Sciences Pvt. Ltd), 20 units RNase inhibitor, and 5 mM DTT in RT buffer (75 mM KCl, 3 mM MgCl_2_, 50 mM Tris, pH 8.4) were added, yielding a 20 μL reaction. The reaction was incubated at 50 °C for 90 min, and 70 °C for 15 min. PCR amplification was then carried out with primers 1 and 2 (Table S2), and PCR products were gel-purified and restriction digested, and cloned and sequenced.

### Sequencing of 3’ ends of cDNAs

The cDNA portions of RNA-cDNA molecules (800 ng) resulting from bRT-Avd activity on DGR RNA were poly-dA tailed by incubation in 200 μL with 40 units of terminal deoxynucleotidyl transferase (TdT, NEB), 0.25 mM CoCl_2_, and 0.2 mM dATP in TdT buffer at 37 °C for 45 min, and 70 °C for 10 min. The sample was P:C:I and C:I extracted, and AA/E-glycogen precipitated. The pellet was re-suspended in water and used for PCR amplification using primers 3 and 4 (Table S2). PCR products were gel-purified, restriction digested, and cloned and sequenced.

Complementary DNAs produced from non-DGR templates were poly-dA tailed, as above, except the reaction volume was 50 μL and 10 units of TdT were used, and the reaction was carried out at 37 °C for 30 min, and 75 °C for 20 min. The sample was treated as above, except that primers 5’-pET28b and 3’-pET28b were used for PCR amplification (Table S2).

### RNA cleavage by bRT-Avd

A 20 μL reaction consisting of 1 μg of [α-^32^P]UTP internally-labeled DGR RNA, 75 mM KCl, 3 mM MgCl_2_, 10 mM DTT, 50 mM HEPES, pH 7.5, 20 units RNase inhibitor (NEB),10 % glycerol was incubated with 1.8 μM of bRT-Avd for 2 h at 37 °C, in the presence or absence of 100 μM dNTPs. In the case of nonhydrolyzeable dCTP, 100 μM 2’-deoxycytidine-5’-μ(α,β)-methyleno]triphosphate (Jena Bioscience) was used.

### 3’-biotinylated DGR RNA

RNA was biotinylated using the Pierce™ RNA 3’ end biotinylation kit, according to manufacturer’s instructions. Reverse transcription with bRT-Avd and 3’-biotinylated RNA was carried out as described above. After proteinase K treatment, 1 μL of the reaction was incubated with 5 μL Dynabeads M-280 (Invitrogen), which had been equilibrated in binding buffer (1 M NaCl, 50 mM Tris, pH 8, 0.1 % Triton X-100, and 5% nuclease free-BSA). The sample was then incubated at RT for 20 min with constant shaking, and separated from solution magnetically. Beads were washed 10x with 1 mL binding buffer, and resuspended in 100 μL of 150 mM NaCl, 50 mM Tris, pH 8. PCR was carried out with 5 μL of the resuspended beads using primers 1 and 5 (Table S2).

### Hybrid molecules

RNA spanning Sp 56-140 (5’ phosphorylated and containing a 3’ dideoxy) with either adenosine or deoxyadenosine at Sp 56 was chemically synthesized (IDT) and gel-purified. Chemically synthesized molecules (12 μM) were incubated with *in vitro* transcribed and gel-purified RNA spanning *avd* 91-Sp 55 or *avd* 368-Sp 55 (3.4 μM) in the presence of 8 μM splint-1 DNA oligonucleotide (Table S2), 8% DMSO, and 0.2x T4 RNA Ligase 1 (T4Rnl1) buffer in 250 μL. The splint was annealed to the RNA molecules by heating at 95 °C for 3 min and cooling at 0.2 °C/min to 20 °C. A 250 μL solution consisting of 0.8x T4Rnl1 buffer, 2 mM ATP, 4% DMSO, and 540 units T4Rnl1 was mixed with the annealed reaction. The reaction was incubated for 5 h at 37 °C. The sample was then extracted with phenol:chloroform (P:C) and AA/E-glycogen precipitated. The pellet was resuspended in water and gel-purified. Hybrid molecules for cleavage experiments were prepared similarly, except the *in vitro* transcribed RNA was internally-labeled with [α-^32^P]UTP.

For detection, hybrid molecules (2 pmol) were treated with Shrimp Alkaline Phosphatase (2 units) in 20 μL cutsmart buffer for 30 min at 37 °C, and 10 min at 65 °C. The sample was purified using G-25 desalting column, and the RNA was 5’ ^32^P-labeled in a 40 μL reaction containing 4 μCi [γ-^32^P] ATP and 20 units of T4 polynucleotide kinase (NEB, PNK) in PNK buffer. The reaction was carried out at 37 °C for 45 min, and 65 °C for 20 min. The sample was purified using a G-25 desalting column.

### Sequencing of RNA cleavage product

Gel-purified cleaved RNA (400 ng), 4 μM adapter DNA, and 5.2 μM splint-2 DNA (Table S2) were annealed in 6% DMSO and 0.2x T4Rnl1 buffer in 25 μL by heating at 95 °C for 3 min and cooling at 0.2 °C/min to 20 °C (Table S2). A 25 μL solution consisting of 0.8x T4Rnl1 buffer, 2 mM ATP, 4% DMSO, and T4Rnl1 (60 units) was mixed with the annealed reaction, and the mixture was incubated for 5 h at 37 °C. The sample was then P:C extracted and AA/E-glycogen precipitated. The pellet was resuspended in water and amplified by RT-PCR using primers 6 and 7 (Table S2). RT-PCR products were gel-purified and restriction digested, and cloned and sequenced.

### RNase Protection Assay and Small RNA Sequencing

A 50 μL reaction consisting of 100 ng of [α-^32^P]UTP internally-labeled RNA, 75 mM KCl, 3 mM MgCl_2_, 10 mM DTT, 50 mM HEPES, pH 7.5, 10 % glycerol was incubated with 1.8 μM of bRT, Avd, or bRT-Avd for 20 min at 37 °C. RNase A/T1 mix (2 μL) was added and the incubation was carried out a further 30 min at 37 °C. The reaction was stopped through P:C:I extraction and AA/E-glycogen precipitation.

For sequencing the protected RNA, the reaction was scaled up five-fold with unlabeled RNA. Following purification, the protected RNA was dephosphorylated using alkaline phosphatase (NEB), and 5’-phosphorylated using T4 polynucleotide kinase (NEB), according to manufacturer’s directions. This 5’-phosphorylated RNA sample was used for cDNA library preparation using the Illumina TruSeq small RNA library preparation kit according to manufacturer’s directions, and sequencing was carried out at the Institute of Genomic Medicine, University of California, San Diego. Illumina small RNA sequencing output reads were trimmed with cutadapt 1.9.0 (28) using the adapter sequence TGGAATTCTCGGGTGCCAAG. Trimmed reads were aligned to a DGR reference sequence (corresponding to the DGR RNA) using bowtie2-align-s version 2.2.6 with default parameters, and unmapped reads were filtered using samtools (29,30). Reads that successfully mapped to the DGR were analyzed with a Python script to determine regions of the DGR sequence that were statistically enriched with mapped reads. Briefly, read start position counts were binned in ~10 basepair regions and tested for binomial enrichment within each bin with Bonferroni correction for the number of bins tested. Reads within each significantly enriched region were assembled through overlap consensus (31).

### Quantification of dNTPs in *Bordetella*

*Bordetella bronchiseptica* ML6401 was grown overnight with shaking at 37 °C in LB supplemented with streptomycin (20 μg/mL) and nicotinic acid (10 mM) to OD_600_ ~0.4. One mL of bacterial culture was centrifuged, and the bacterial pellet was washed three times with 1 mL of ice cold PBS, centrifuged again and resuspended in 70% CH_3_OH. Internal standards (50 pmol ^13^C^15^N-dATP, 100 pmol ^13^C^15^N-dCTP, 50 pmol ^13^C^15^N-dGTP, 50 pmol ^13^C^15^N-TTP) were added to the samples, and the samples were stored overnight at −80 °C. Samples were then thawed and centrifuged, and the supernatant was air-dried and stored at −20 °C. Samples were reconstituted in 200 μL 2 mM NH_3_H_2_PO_4_ with 3 mM hexylamine, and 20 μL were injected for dNTP analysis by liquid chromatography-tandem mass spectrometry (LC-MS/MS). Intracellular dATP, dCTP, dGTP, TTP and dUTP were separated by Ultimate 3000 LC system (Thermo Scientific) using a Kinetex EVO C18 column (Phenomenex). An API5000 triple quadrupole mass spectrometer (ABsciex, MA, USA) was used for detection by multiple reaction monitoring. For estimation of nucleotide concentrations, a single cell volume of 4.5 fL determined for *E. coli* was used (32), based on the similarity between *B. bronchiseptica* and *E. coli*, both rod-shaped bacteria.

## RESULTS

### Reconstitution of cDNA synthesis

The bRT-Avd complex was co-expressed and co-purified, in apparent 1:5 stoichiometry, and incubated with *in vitro* transcribed and gel-purified DGR RNA (Figs. 1B and S1B, C). This ~580 nucleotide (nt) RNA contained *TR* (134 nt), and extensive upstream (297 nt from *avd*) and downstream sequences (144 nt of spacer between *TR* and *brt*, and 6 nt of *brt*). The extensive *avd* sequence was based on prior results showing that 300 nt upstream of *TR* are required for near wild-type retrohoming activity (23). Shorter lengths of *avd*, especially below 115 nt, lead to ~500-1400-fold lower retrohoming activity. The downstream sequence was based on studies described below showing that nearly the entire length of the spacer region was required for cDNA synthesis. In reverse transcription reactions carried out in the absence of an exogenous primer and monitored through the incorporation of [α-^32^P]dCTP, we observed production of a major radiolabeled band with an apparent size similar to the template RNA, ~580 nt (Figs. 1C and S1D, E). This 580 nt band, however, was not entirely composed of cDNA, as it was digested by RNase into smaller bands, primarily ~120 and ~90 nt in length (both slightly smaller than the 134-nt *TR*), along with a number of minor bands (Figs. 1C and S1F, G). These 120-and 90-nt bands corresponded to cDNAs as indicated by their sensitivity to DNase. All bands were sensitive to combined RNase and DNase digestion.

Evidence that these cDNAs corresponded to *TR*-cDNA came from templates deleted internally within *TR* (Fig. 1D). A template containing an internal 40 nt deletion in *TR* resulted in cDNA products that were correspondingly 40 nt shorter than the 120 nt and 90 nt cDNAs, that is, ~80 and ~50 nt (Fig. 1E). A template containing an internal 80 nt deletion in *TR* resulted in a single cDNA product which was correspondingly 80 nt shorter than the 120 nt cDNA (Fig. 1E). A second shorter cDNA was not observed, suggesting that the reverse transcription termination site that leads to the 90 nt cDNA was missing in the *TR*20-Δ80 construct. This supposition was confirmed by sequencing of cDNAs, as detailed below. A template containing an internal 100 nt deletion in *TR* resulted in a single weak cDNA product which was correspondingly 100 nt shorter than the 120 nt cDNA (Fig. 1E). These results provided evidence for template-primed reverse transcription, which resulted in *TR*-cDNA molecules covalently linked to the template RNA, as observed *in vivo* (19).

Curiously, the 580 nt radiolabeled band resulting from reverse transcription was sensitive to RNase but did not appreciably change size upon DNase digestion (Fig. 1C). To address this, we analyzed the 580 nt band on a low percentage gel. Several distinct species were resolved — a high-intensity ~580 nt species (Fig. 1F, red arrowhead) and several lower-intensity and smaller (~540 nt) species (Fig. 1F, green arrowheads). These smaller species were also visible upon overloading (Fig. S1D). The nature of these species was clarified by blocking the 3’-OH of the RNA through periodate treatment prior to reverse transcription (33,34). Most significantly, the amount of the 580 nt species was substantially diminished upon periodate-treatment, but the yields of the 90 and 120 nt *TR*-cDNAs were unchanged (Fig. 1F). These data indicate that the 580 nt species lacked *TR*-cDNAs, and instead consisted of the RNA template with one or just a few dNTPs added to its 3’ end in nontemplated fashion (3’ addition activity). DNase does not degrade further than tetranucleotides on average (35), explaining why the radiolabel on the 580 nt species survives DNase treatment. These data further suggest that covalently linked RNA-cDNA molecules were found in the several 540 nt species. The nature of these RNA-cDNA molecules, which migrate faster than the input 580 nt RNA template, is investigated below. bRT alone produced the 580 nt band but no cDNA (Figs. 1C and S1G), indicating that it had the presumably artifactual 3’ addition activity but no reverse transcription activity. In support of this conclusion, when bRT alone was reacted with periodate-treated RNA, no radiolabeled products were formed (Fig. S1H). No radiolabeled products resulted from Avd alone, demonstrating that Avd had neither cDNA synthesis nor 3’ addition activity (Figs. 1C and S1G).

### Adenine-mutagenized *TR*-cDNA

The sequence of cDNAs produced by bRT-Avd were identified from RNA-cDNA molecules that were amplified by RT-PCR (Fig. 2A, Table S2). Amplicons were evident specifically for templates reacted with bRT-Avd, but not bRT or Avd alone, or no protein (Fig. 2B). Sequencing of these amplicons showed that they contained *TR*, and remarkably ~50% of *TR* adenines were substituted (Figs. 2C and S2), identical to the ~50% observed *in vivo* (1,4). Substitutions at other bases were detected, but at extremely low frequency (0-0.5%), as also reported *in vivo* (1), and rare single basepair deletions and insertions could be detected as well (0.1%). Adenine-specific misincorporation by bRT-Avd was confirmed by its ability to synthesize *TR*-cDNA in the absence of TTP, but not dATP or dGTP (Figs. 2D and S3); dCTP was not omitted, as [α-^32^P]dCTP was used for radiolabeling. Synthesis excluding TTP resulted in greater production of the 90 nt product as compared to the 120 nt product and exhibited slower kinetics, most likely because misincorporation was required at every *TR* adenine.

**Figure 2.**
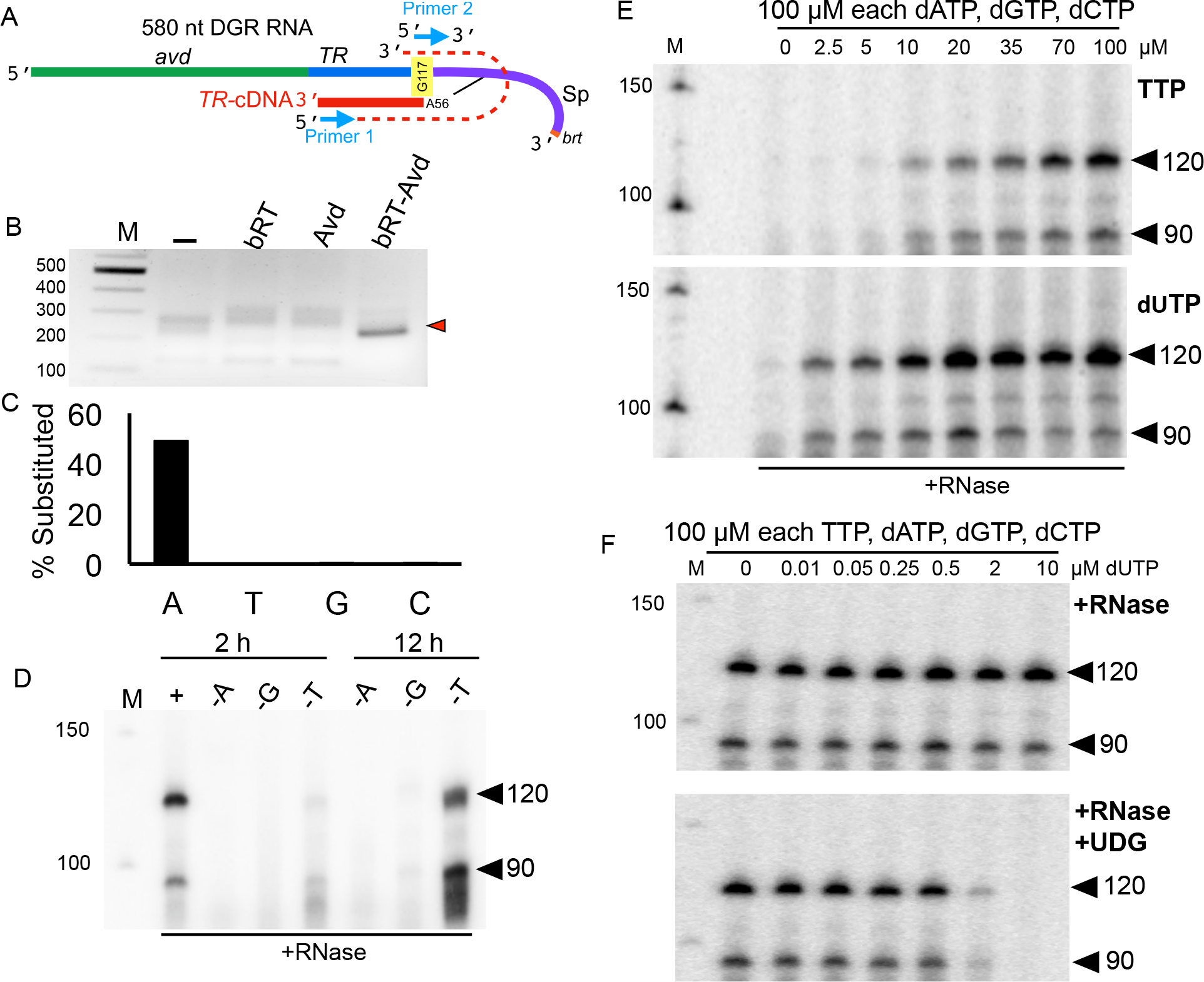
Adenine Mutagenesis and Template-Priming. (A) Covalently-linked RNA-cDNA molecule. The linkage is to Sp A56 of the RNA, and the first nucleotide reverse transcribed is *TR* G117. The RT-PCR product resulting from primers 1 and 2 (blue arrows) is indicated by the dashed red line. (B) RT-PCR amplicons from 580 nt DGR RNA reacted with no protein (-), bRT, Avd, or bRT-Avd, separated on a 2% agarose gel and ethidium bromide-stained. The specific amplicon produced from reaction with bRT-Avd shown by the red arrowhead. (C) Percentage of substitutions in *TR*-cDNA determined by sequencing. (D) Radiolabeled 120 and 90 nt cDNA products, indicated by arrowheads, resulting from bRT-Avd activity with the 580 nt DGR RNA as template for 2 h (left) or 12 h (right). Either standard dNTPs (dATP, dGTP, dCTP, TTP), as indicated by “+”,were present in the reaction, or standard dNTPs excluding dATP (-A), dGTP (-G), or TTP (-T) were present. Products were treated with RNase, and resolved by denaturing PAGE. (E) Radiolabeled 120 and 90 nt cDNA products, indicated by arrowheads, resulting from bRT-Avd activity for 2 h with the 580 nt DGR RNA as template with varying TTP (top) or dUTP (bottom) concentrations. Products were treated with RNase, and resolved by denaturing PAGE. (F) Radiolabeled 120 and 90 nt cDNA products, indicated by arrowheads, resulting from bRT-Avd activity for 2 h with the 580 nt DGR RNA as template with varying dUTP concentrations. Products were either RNase-treated (top), or both RNase-and UDG-treated (bottom), and resolved by denaturing PAGE

We also found that bRT-Avd was able to incorporate dUTP, but not UTP or dUMP, into *TR*-cDNA (Figs. S4A, B). Incorporation was assessed by scission of uracylated DNA upon digestion with uracil DNA glycosylase (UDG) and heating (36). bRT-Avd had an apparent ~10-fold preference for dUTP over TTP (Figs. 2E and S4C), and incorporated dUTP even when the dUTP concentration was 50-fold lower than that of competing dNTPs (Figs. 2F and S4D). As validation of these assays, HIV-1 RT was shown to have a marginal preference for TTP over dUTP, as previously reported (37) (Figs. S4E-G). In *Bordetella*, however, dUTP was measured to be >200-fold lower in concentration than TTP (Table S3), indicating that mutagenesis *in vivo* almost certainly proceeds through misincorporation of dATP, dGTP, or dCTP at template adenines rather than incorporation of dUTP followed by de-uracylation and repair (1, 4).

Together, these results establish that bRT-Avd is sufficient for template-primed *TR*-cDNA synthesis from DGR RNA, and that bRT-Avd displays a vigorous promiscuity at template adenines but fidelity at other bases.

### Priming from Sp 56

Sequencing of RT-PCR amplicons generated from RNA-cDNA molecules showed that immediately adjacent to *TR* was sequence from the noncoding spacer (Sp) region (Figs. 1A and S2). *TR* and Sp sequences originated from different strands (i.e., sense vs. antisense), signifying that Sp had folded back onto *TR* during priming (Fig. 3A). The precise site of priming was ambiguous due to sequence degeneracy, but was clarified by the pattern of RNA substitutions in *TR* or Sp transferred to RNA-cDNA molecules (Figs. S5A and S5B). In short, we found that substitutions at Sp 55 and 56, but not *TR* 118 and 119, were transferred (Fig. S5C). These data showed that synthesis was primed from Sp A56, and that *TR* G117 was the first nucleotide reverse transcribed (Fig. S5D). *TR* G117 is located in a GC-rich region of the *TR* that is not prone to adenine-mutagenesis. These assignments are identical to those derived from analysis of RNA-cDNA molecules produced *in vivo* (19).

**Figure 3.**
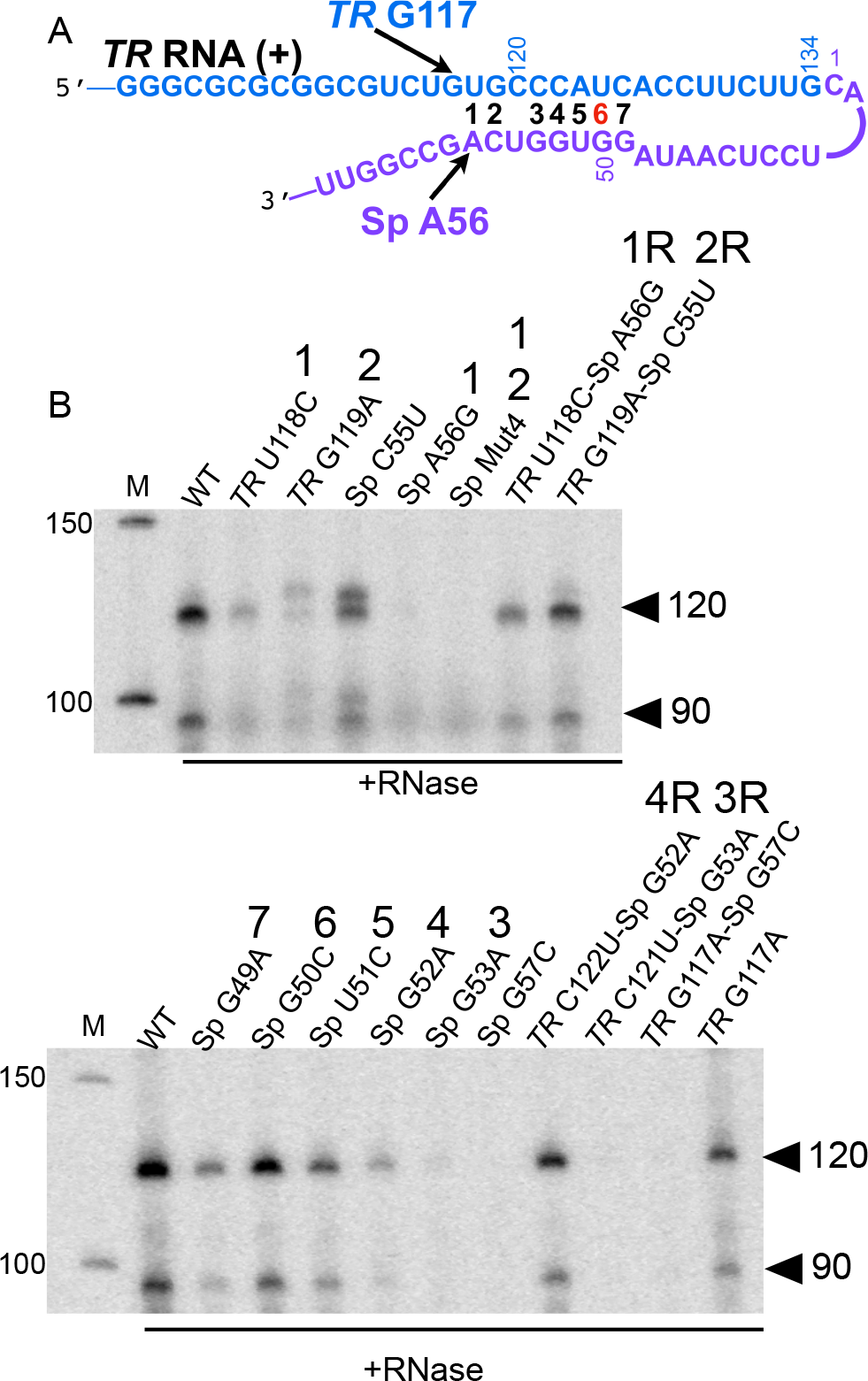
*TR*-Sp Interactions. (A) Complementarity between *TR* (blue) and Sp (purple) segments. Potential basepairs are numbered (wobble in red). (B) Radiolabeled 120 and 90 nt cDNA products, indicated by arrowheads, resulting from bRT-Avd activity for 2h with the WT or mutated 580 nt DGR RNA as template. Products were treated with RNase, and resolved by denaturing PAGE. Numbers over lane labels correspond to the basepair tested by the mutation, with “R” referring to restoration of the basepair. Sp Mut4 corresponds to Sp 55-CAGC substituted with 55-GUCG

The 3’ ends of cDNAs were identified by poly-dA tailing and PCR amplification (Fig. S6A). Amplicons were once again evident specifically from templates reacted with bRT-Avd, but not with bRT or Avd alone, or no protein (Fig. S6B). Three length classes of cDNAs, all adenine-mutagenized, were identified (Fig. S6C). The longest cDNAs were 118-122 nt, having terminated within the six 3’ bases of *avd*; these corresponded to the most abundant radiolabeled species, the 120 nt band (Figs. 1C, S1F). The second longest were almost all 94-96 nt, having terminated between *TR* 22-24; these corresponded to the second most abundant radiolabeled species, the 90 nt band. Notably, codons permissive to adenine mutagenesis occur exclusively between *TR* 23-98, and thus both the 90 and 120 nt cDNAs carry the full potential of adenine-mutagenesis. Short cDNAs of 40-59 nt, for which synthesis had terminated between *TR* 59-78, were also found, and these corresponded to a minor population of radiolabeled cDNAs that became evident only after prolonged reaction times (Fig. S3). The relative proportion of these cDNAs did not change over time, indicating independent termination events rather than precursor-product relationships. The internally deleted *TR*20-Δ80 and *TR*10-Δ100 DGR RNAs (Fig. 1D) lack the *TR* 22-24 termination site, explaining why they produced one and not two cDNA products.

### *TR*-Sp Interactions

Seven basepairs (bp), including a wobble, can be formed between sequences proximal to the initiation site in *TR* and the priming site in Sp (Fig. 3A). Basepairing appeared to be important at three of these seven (bp 1, 2, and 4), as disruption led to greatly attenuated cDNA synthesis, and complementary substitutions rescued cDNA synthesis (Figs. 3B and S7). For the complementary substitution at bp 1 (*TR* U118C-Sp A56G), Sp 56G was the priming nucleotide (Fig. S5C), indicating that the identity of the base at Sp 56 was not strictly essential to priming. For bp 2, which is adjacent to the priming site, disruption resulted in additional cDNA bands, which were due to non-templated insertion of four to seven nucleotides (Fig. S5C). Disruption of bp 3 caused a drastic reduction in cDNA synthesis, but a complementary substitution was ineffective at restoring synthesis, suggesting that guanine is required at Sp 53. Basepairs 5, 6, and 7 appeared not to be crucial, as disruptions affected cDNA synthesis only mildly. The *in vitro* results for bp 1 and 2 are similar to those found *in vivo* (19), including the addition of non-templated nucleotides. The other potential basepairs were not examined *in vivo*.

We asked whether the priming site could be shifted by introducing an additional basepair downstream of Sp 56, at Sp 57 (Sp G57C-*TR* G117). However, this nearly abrogated cDNA synthesis (Figs. 3B and S7). Disruption of this artificial basepair (Sp G57C-*TR* G117A) did not rescue cDNA synthesis (*TR* G117A by itself was competent for cDNA synthesis), indicating a specific role for guanine at Sp 57.

In sum, these results provide evidence for limited but not extensive basepairing between *TR* and Sp, and also suggest higher-order organization of the template, with specific bases required at certain positions.

### RNA-cDNA Linkage

The apparent 580 nt size of the radiolabeled band produced by bRT-Avd was puzzling, given a 580 nt RNA template and 90 or 120 nt cDNAs. Two possibilities were examined. The first was that the RNA had been cleaved at Sp 56, thereby freeing its 3’-OH for priming. We first asked whether a template truncated at Sp 56 was competent for cDNA synthesis, and found that it was not (Fig. S8). This result indicated that regions downstream of Sp 56 were essential for cDNA synthesis, but did not rule out cleavage at Sp 56. Cleavage was directly examined with an internally-labeled 580 nt DGR RNA incubated with bRT-Avd in the presence or absence of dNTPs. No cleavage products were detected, with as much as 10-fold overloading of the RNA (Fig. S9).

To test the internal priming hypothesis, we biotinylated the 3’ end of the 580 nt DGR RNA, and used that as a template for reverse transcription by bRT-Avd (Fig. 4A). Following reverse transcription, 3’-biotinylated RNA was isolated using streptavidin beads, and the presence of *TR*-cDNA accompanying the intact RNA was probed by PCR. A specific PCR amplicon of the correct size was evident for 3’-biotinylated RNA reacted with bRT-Avd, but not for 3’-biotinylated RNA reacted with no protein or unbiotinylated RNA reacted with bRT-Avd (Fig. 4B). The PCR amplicon was confirmed to correspond to adenine-mutagenized *TR*-cDNA by sequencing (Fig. S10A), and input samples were confirmed to all contain RNA by RT-PCR (Fig. S10B); the two input samples reacted with bRT-Avd to contain *TR*-cDNA by PCR (Fig. S10C); and the two purified biotinylated samples to contain RNA by RT-PCR (Fig. S10B). We also verified that the RNA-cDNA molecules examined were in *cis* (part of the same molecule) and not in *trans* (two or more RNA molecules associated through a common *TR*-cDNA or other means). This was done by carrying out the reverse transcription reaction with unbiotinylated RNA, inactivating bRT-Avd with proteinase K as in all experiments, and then adding an equal quantity of unreacted biotinylated RNA prior to isolation with streptavidin beads. We found no recruitment of unbiotinylated RNA-cDNA molecules by biotinylated RNA (Fig. S10D), demonstrating that the molecules examined were indeed in *cis.* These results indicated that the streptavidin beads had effectively and specifically captured the biotin at the 3’ end of the RNA, and that the captured and intact RNA contained *TR*-cDNA. This result also showed that internally located Sp 56 can serve as a priming site, with its 2’-OH being the most chemically plausible site for priming. RNA-cDNA linkage through a 2’-OH has been observed previously in retrons (38–41), in which a specific guanosine internally located in an RNA template primes cDNA synthesis by the retron reverse transcriptase, resulting in a branched RNA-cDNA molecule linked by a 2’-5’ phosphodiester bond.

**Figure 4.**
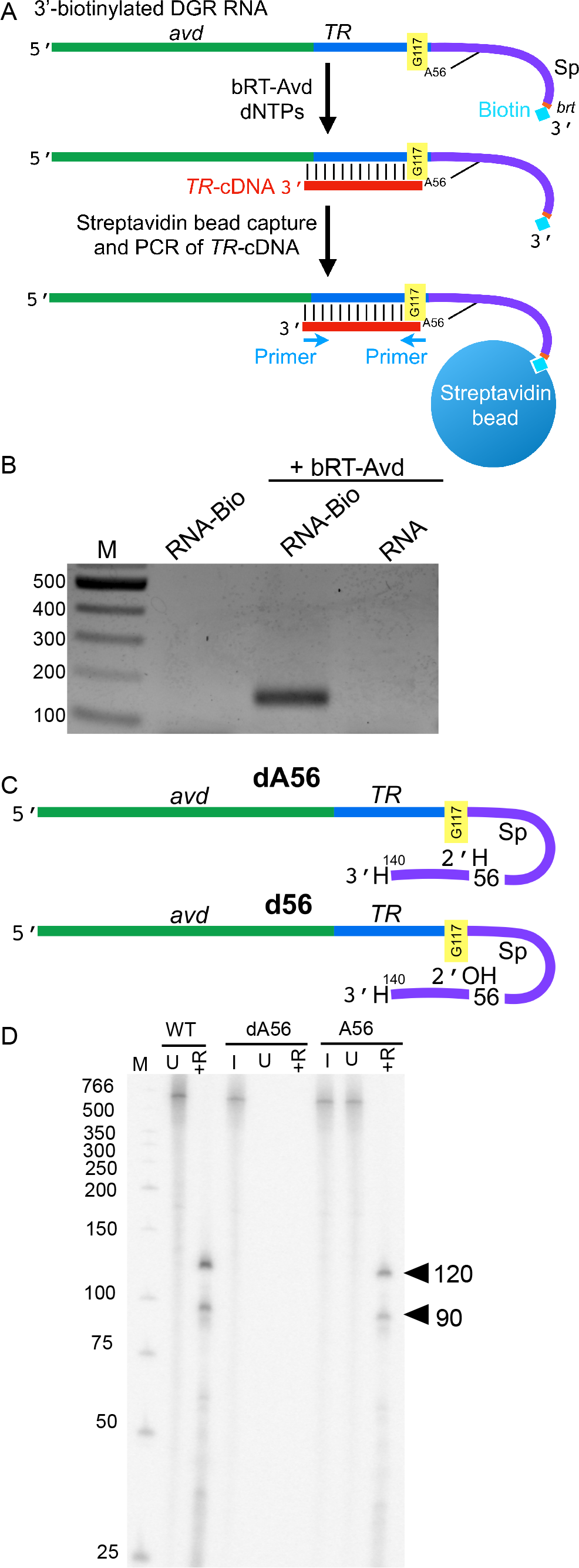
Branched RNA-cDNA. (A) The 580 nt DGR RNA was biotinylated at its 3’ end and used as a template for reverse-transcription by bRT-Avd, after which biotinylated RNA was captured with streptavidin beads, and the presence of *TR*-cDNA was detected by PCR using the indicated primers. (B) The 580 nt DGR RNA was biotinylated at its 3’ end (RNA-Bio), and either reacted with no protein or used as a template for reverse transcription with bRT-Avd. The 580 nt DGR RNA in its unbiotinylated form (RNA) was also used as a template for reverse transcription with bRT-Avd. Samples were then purified using streptavidin beads, and the presence of *TR*-cDNA in the purified samples was assessed by PCR. Products from the PCR reaction were resolved on an agarose gel. (C) Hybrid dA56 580 nt DGR RNA containing deoxyadenosine at Sp 56 (indicated with H at 2’ position) and hybrid d56 580 nt DGR RNA containing adenosine at Sp 56 (indicated with OH at 2’). Both molecules terminate at Sp 140 and have a dideoxynucleotide at the 3’ end (indicated with H at 3’). (D) Radiolabeled products resulting from bRT-Avd activity for 12 h with 580 nt DGR RNA, hybrid 580 nt dA56, or hybrid 580 nt A56 DGR RNA as template. Products were untreated (U) or RNase-treated (+R), and resolved by denaturing PAGE. Separate samples of dA56 and A56 were 5’ ^32^P-labeled for visualization of input templates (I). The positions of the 120 and 90 nt cDNAs are indicated

To test the importance of the 2’-OH of Sp 56 in priming, we used chemical synthesis and ligation to create a hybrid 580 nt DGR RNA with a deoxyadenosine at Sp 56 (dA56), and as a control, an equivalent molecule with ribonucleotides throughout (A56) (Fig. 4C). For the efficiency of ligation, it was necessary to use a template that ended at Sp 140 and had a 3’ dideoxynucleotide. The latter precluded the 3’ addition activity of bRT. The A56 molecule supported cDNA synthesis, albeit less efficiently than fully *in vitro* transcribed DGR RNA (48%), which was likewise truncated at Sp 140 (Fig. 4D). In contrast, an equivalent quantity of the dA56 hybrid molecule was incompetent for cDNA synthesis, demonstrating that the 2’-OH of Sp 56 was essential for priming. Taken together, these results support the conclusion that priming from the 580 nt DGR RNA occurs from Sp 56 without RNA cleavage, resulting in branched molecules linked by 2’-5’ RNA-cDNA phosphodiester bonds. Such RNA-cDNA branched molecules apparently septate through gels faster than the unreacted RNA template, as shown by the ~540 nt species corresponding to RNA-cDNA molecules (Fig. 1F).

While no RNA cleavage was evident for the 580 nt DGR RNA template, we did detect cleavage of a shorter template, called the ‘core DGR RNA’. This 294-nt template had just 20 nt of *avd* at its 5’ end and extended to Sp 140 at its 3’ end (Fig. 5A). The core DGR RNA supported cDNA synthesis (Fig. 5B), whereas a template lacking the entire *avd* sequence at the 5’ end or the entire Sp sequence at the 3’ end did not (Figs. S11A and B). Similarly, templates that contained only 10 nt of *avd* at the 5’ end or were deleted of 20 nt of Sp at the 3’ end did not support cDNA synthesis (Fig. S11C). These experiments showed that the extensive 300 nt sequence of *avd* upstream of *TR* required for near wild-type retrohoming activity (23) was almost entirely dispensable for cDNA synthesis. Notably, reverse transcription from the core DGR RNA resulted in ~7-fold greater cDNA production (i.e., the 90 and 120 nt cDNAs) as compared to the 580 nt DGR RNA (Figs. S11C and S12A). We verified that cDNAs produced from the core DGR RNA were still primed from Sp 56 with *TR* G117 the first nucleotide reverse transcribed through sequencing of amplified RNA-cDNA molecules (Figs. S12B and C).

**Figure 5.**
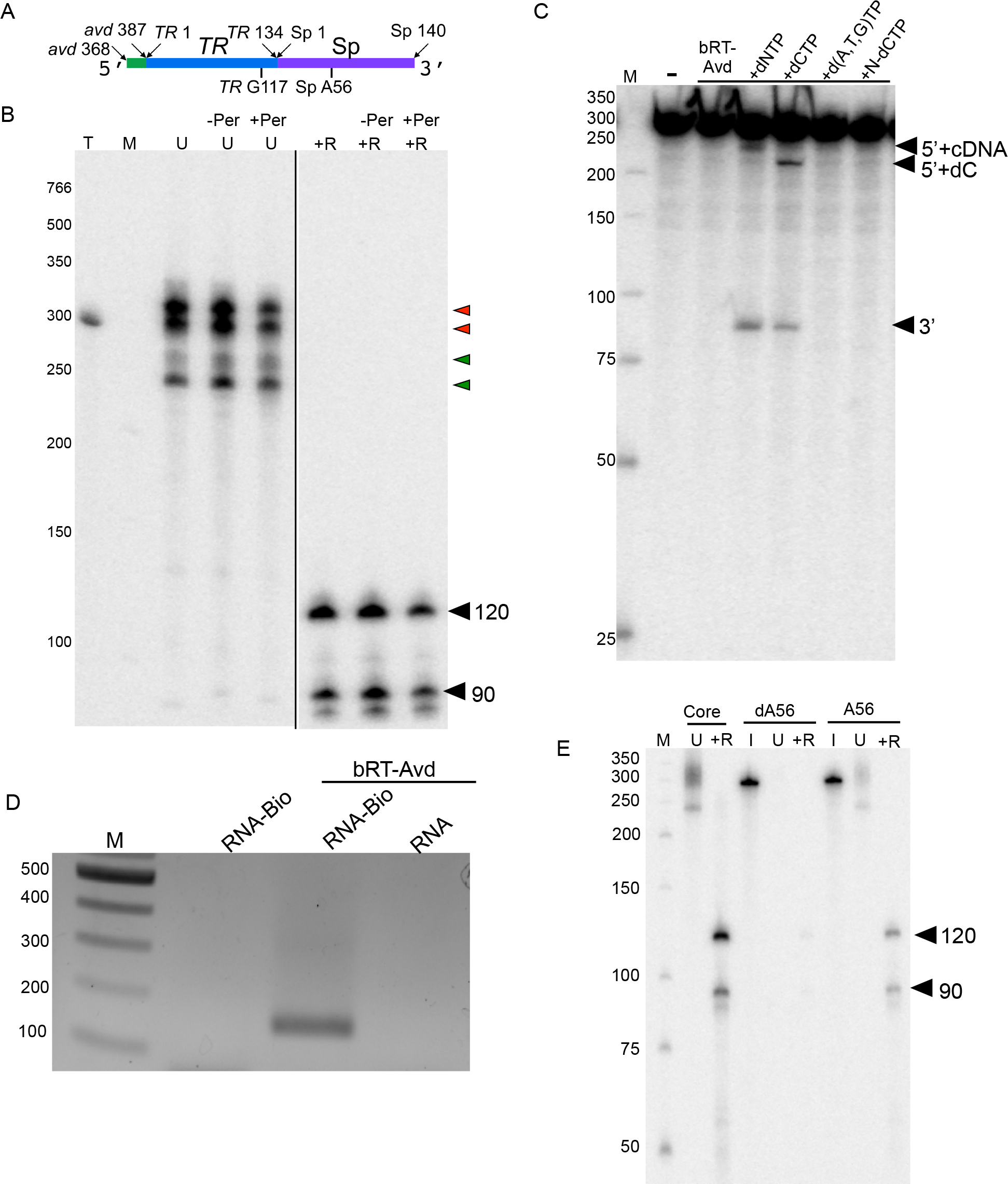
Core DGR RNA. (A) Schematic of core DGR RNA. (B) Radiolabeled products resulting from bRT-Avd activity for 2 h with the core DGR RNA as template. Prior to the reverse transcription reaction, the RNA template was untreated (-Per) or treated with periodate (+Per). Products from the reaction were untreated (U) or treated with RNase (+R), and resolved by 6% denaturing PAGE. Lane T corresponds to internally-labeled core DGR RNA as a marker for the size of the template. Red arrowheads indicate radiolabeled product bands that migrate at the same position or slower than the core DGR RNA, and green arrowheads ones that migrate faster. The positions of the 120 and 90 nt cDNA bands are indicated. The two panels are from the same gel, with the black line indicating that intermediate lanes were removed. (C) Internally-labeled core DGR RNA was not incubated (−), or incubated with bRT-Avd alone or bRT-Avd with 100 μM standard dNTPs (+dNTP), 100 μM dCTP (+CTP), 100 μM dNTPs excluding dCTP (+d(A,T,G)TP), or 100 μM nonhydrolyzeable analog of dCTP (+N-dCTP) for 2 h. Incubation products were resolved by denaturing PAGE. The band corresponding to the 5’ fragment of the cleaved core RNA containing either a deoxycytidine alone (5’+dC) or cDNA (5’+cDNA), and the band corresponding to the 3’ fragment of the RNA are indicated. (D) The core DGR RNA was biotinylated at its 3’ end (RNA-Bio), and either reacted with no protein or used as a template for reverse transcription with bRT-Avd. The core DGR RNA in its unbiotinylated form (RNA) was also used as a template for reverse transcription with bRT-Avd. Samples were then purified using streptavidin beads, and the presence of *TR*-cDNA in the purified samples was assessed by PCR. Products from the PCR reaction were resolved on an agarose gel. (E) Radiolabeled products resulting from bRT-Avd activity for 12 h with core, hybrid core dA56, or hybrid core A56 DGR RNA as template. Products were untreated (U) or treated with RNase (+R), and resolved by denaturing PAGE. Separate samples of core dA56 and A56 were 5’ ^32^P-labeled for visualization of inputs (I). The positions of the 120 and 90 nt cDNAs are indicated

Most interestingly, several RNA-cDNA bands were produced from the core DGR RNA, with one of these having a mobility greater than the unreacted template (Fig. 5B). To delve into these RNA-cDNA molecules further, we asked whether the core DGR RNA had been cleaved. Upon incubation with bRT-Avd, cleavage of ~10% of internally-labeled core DGR RNA was evident, but only in the presence of dNTPs (Fig. 5C). Indeed, the initiating dNTP (i.e., dCTP, which basepairs to *TR* G117) was necessary and sufficient for cleavage, and hydrolysis of dCTP was required, as cleavage did not occur with a nonhydrolyzeable dCTP analog. Since the core and 580 nt DGR RNAs differed at both 5’ and 3’ ends, we examined cleavage of a 580 nt DGR RNA lacking just the 3’ end (3’Δ10). Cleavage was not evident (Fig. S12D), indicating that the length of the *avd* sequence at the 5’ end modulated cleavage.

The sizes of the cleavage products were consistent with bond breakage circa Sp 56. The larger cleavage product was ~210 nt in length when dCTP was added and >210 nt when dNTPs were added, indicating that it included cDNA linked to Sp 56 and corresponded to the 5’ end of the core DGR RNA. The smaller cleavage product had a size of ~80 nt regardless of whether dNTPs or dCTP alone were added, indicating that it did not include cDNA or the Sp 56 priming site, and corresponded to the 3’ end of the core DGR RNA. The 3’ cleavage product was purified, amplified by RT-PCR following ligation of an adaptor to its 5’ end, and cloned and sequenced. This showed that, surprisingly, Sp 58 and not Sp 57 was at its 5’ end, indicating that cleavage had occurred between Sp 57-58 (Figs. S12E and F). The 3’ cleavage product was also phosphorylated at its 5’ end prior to adaptor ligation to account for the possibility that some cleaved molecules may have lacked a 5’ phosphate. The sequencing results from this experiment were identical, showing that Sp 58 was at the 5’ end. We attempted a parallel method for sequencing the 3’ end of the larger cleavage product, but were unsuccessful, perhaps due to deoxycytidine at Sp 56 blocking adapter ligation. We conclude from these data that synthesis-dependent cleavage occurs between Sp 57-58, but cannot rule out the possibility that a second cleavage event occurs between Sp 56-57.

To determine whether cleavage was necessary for cDNA synthesis from the core DGR RNA, we used 3’-biotinylated core DGR RNA as a template for reverse transcription, as described above (Fig. 4A). We once again captured the 3’-biotinylated RNA following reverse transcription with streptavidin beads, and asked whether *TR*-cDNA accompanying the intact RNA was present by PCR. We observed a specific amplicon of the correct size for 3’-biotinylated core DGR RNA reacted with bRT-Avd, but not for 3’-biotinylated RNA reacted with no protein or unbiotinylated RNA reacted with bRT-Avd (Fig. 5D). The PCR amplicon was verified to be adenine-mutagenized *TR*-cDNA by sequencing (Fig. S13A), and the integrity of input and purified samples was verified, as described above for the 580 nt DGR RNA (Figs. S13B and S13C). We also confirmed that RNA-cDNA molecules were in *cis* (Fig. S13D). This result indicated that cleavage was not required for priming, as internally located Sp 56 could serve as the priming site. To verify the importance of the 2’-OH of Sp A56 for priming in the core DGR RNA, we created a hybrid molecule with dA56 and a control molecule with A56, as described above (Fig. 4C). The A56 form of the core DGR RNA supported cDNA synthesis (50% of that from fully *in vitro* transcribed RNA). However, cDNA synthesis was severely attenuated from an equivalent quantity of the core dA56 hybrid molecule (Fig. 5E), showing that the 2’-OH of Sp A56 was crucial for cDNA synthesis from the core DGR RNA. The 2’-OH of Sp A56 was also crucial for cleavage of the core DGR RNA, as the core A56 molecule was cleaved in the presence of bRT-Avd and dCTP, whereas dA56 was not (Fig. S12G). In this experiment, only the *in vitro* transcribed portion of the hybrid molecule was internally-labeled, explaining why only the larger of the two cleavage bands was visible for A56.

With these findings in hand, we sought to assign the RNA-cDNA bands from the core DGR RNA. To this end, the core DGR RNA was treated with periodate prior to reverse transcription to minimize the 3’ addition activity of bRT. Four RNA-cDNA bands were observed (Fig. 5C). The very uppermost band (Fig. 5C, top red arrowhead), at an apparent size of ~330 nt, likely contained the 120 nt cDNA covalently linked to the ~210 nt RNA that had been cleaved at Sp 57, and the next uppermost band, at an apparent size of ~300 nt, likely contained the 90 nt cDNA linked to the cleaved RNA. The two lower bands (Fig. 5C, green arrowheads), with greater mobility than the unreacted core DGR RNA, likely corresponded to branched RNA-cDNA molecules, as seen with the 580 nt DGR RNA.

These results indicate that the core DGR RNA can be cleaved by bRT-Avd and that cleavage is coupled to cDNA synthesis. However, cleavage was not required for cDNA synthesis. The 2’-OH of internally located Sp 56 could serve as the priming site, leading to branched RNA-cDNA molecules.

### Specificity to DGR RNA

We asked whether template-primed cDNA synthesis could also occur on a heterologous template. Using a ~200 nt non-DGR template and no primer, we observed no cDNA synthesis and only 3’ addition activity (Fig. S14A). However, with the addition of an oligodeoxynucleotide (ODN) primer, bRT-Avd synthesized cDNAs, which were as expected not covalently attached to the RNA, but were quite short (~5-35 nt).

We next asked whether template-priming was required for cDNA synthesis from the DGR RNA. When synthesis was primed with an ODN such that *TR* G117 was the first nucleotide reverse transcribed (Figs. S14B and C, P^G117^ primer), a 120 nt *TR*-cDNA and other extended *TR*-cDNAs were produced, along with some shorter ones. This ODN primer inhibited template-primed cDNA synthesis. In contrast, when synthesis was primed by ODNs that annealed to other locations on the DGR RNA, exclusively short ODN-primed cDNAs were produced, just as with the non-DGR template. Important insight into the organization of the DGR RNA template came from the observation that truncation of the DGR RNA at Sp 56 eliminated the synthesis of extended cDNAs, even when synthesis was primed with an ODN that annealed to *TR* and primed synthesis from *TR* G117. Only short cDNAs were produced from the truncated template (Fig. 6A, P^G117^ primer). Deletion of the *avd* sequence at the 5’ end of the 580 nt DGR RNA greatly reduced but did not eliminate synthesis of extended cDNAs from primer P^G117^ (Fig. S14D), indicating that the *avd* sequence has a role in processive polymerization but a less major one than the Sp region. These results suggest that *avd* and Sp are organized such that *TR* G117 exists at a functionally privileged location that promotes the processivity of bRT-Avd and enables synthesis of extended cDNAs. In addition, the results show that processivity from *TR* 117 is independent of the mechanism of priming, occurring in the case of both template-and ODN-priming.

**Figure 6.**
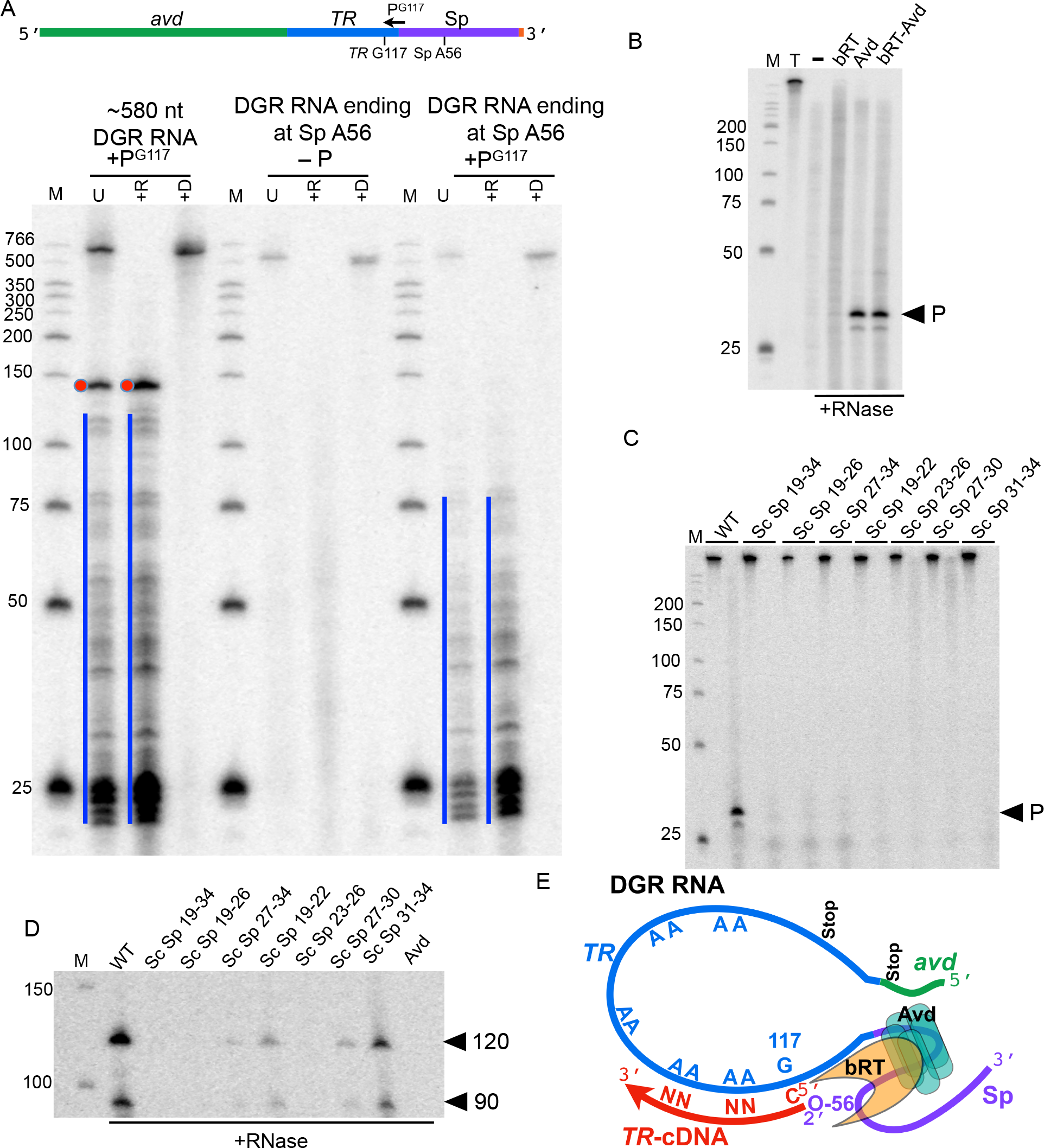
Specificity to DGR RNA. (A) Top, Schematic of DGR RNA template and primer P^G117^. Bottom, radiolabeled products resulting from bRT-Avd activity for 2 h with intact 580 nt DGR RNA or DGR RNA truncated at Sp A56 as template. Reverse transcription reactions were carried out in the absence (-P) or presence of primer P^G117^. Reaction products were untreated (U), treated with RNase (+R), or treated with DNase (+D), and resolved by denaturing PAGE. The blue line indicates ODN-primed cDNA products. The red dot indicates ODN-primed 120 nt cDNA (cDNA + 20 nt primer for a 140 nt band). (B) Protection of internally-labeled 580 nt DGR RNA from RNase by bRT, Avd, or bRT-Avd, with products resolved by 15% denaturing PAGE. The protected band (P) is indicated. (C) RNase protection by Avd, as in panel B, carried out on internally-labeled wild-type 580 nt DGR RNA or 580 nt DGR RNA with scrambled (Sc) Sp sequences, with the first lane in each pair untreated and the second RNase-treated. Products were resolved by denaturing PAGE. The protected band (P) is indicated. (D) Radiolabeled products resulting from bRT-Avd activity for 2 h with the WT 580 nt DGR RNA or DGR RNA containing scrambled (Sc) Sp sequences as template. The last lane shows the activity of Avd alone for 2 h with the WT 580 nt DGR RNA as template. Products were treated with RNase, and resolved by denaturing PAGE. The positions of the 120 and 90 nt cDNAs indicated. (E) Model of processive polymerization of adenine-mutagenized cDNA by bRT-Avd/RNA ribonucleoprotein particle. The 2’-OH of Sp 56 serves as the priming site and forms a 2’-5’ phosphodiester bond with the cDNA. The first nucleotide reverse transcribed is *TR* 117. Adenines in *TR* are unfaithfully reverse transcribed by bRT-Avd (represented by “N”). The RNP promotes processive polymerization, which terminates at one of two stops in the DGR RNA

Both bRT and Avd were required for ODN-primed cDNA synthesis from DGR and non-DGR templates, suggesting a role for Avd in catalysis. Sequencing of short ODN-primed cDNAs synthesized from the non-DGR RNA template revealed adenine-mutagenesis (Fig. S14E). This promiscuity extended to the ability of bRT-Avd to incorporate dUTP into cDNAs synthesized from a non-DGR template (Fig. S14F). These results indicate that adenine-mutagenesis, unlike synthesis of extended cDNAs, is not restricted by the RNA template and is instead intrinsic to bRT-Avd.

To determine the basis for this restriction to the DGR RNA, an RNase protection assay was carried out with internally-labeled 580 nt DGR RNA. A ~30 nt band was protected by Avd and bRT-Avd, but not bRT alone (Fig. 6B). Deep sequencing revealed that this band contained Sp 19-34 (Fig. S15A). The region downstream of Sp 56 was not required for protection by Avd (Fig. S15B). Scrambling the sequence of Sp 19-34, either in its entirety or as 4-or 8-nt segments, resulted in loss of RNase protection by Avd and bRT-Avd (Figs. 6C and S15C, Table S1). Synthesis of cDNA from these templates was also severely attenuated (Figs. 6D and S15C). The exception was scrambled Sp 31-34, for which RNase protection was lost but cDNA synthesis maintained at one-third of wild-type. Apparently, Avd binding to this template was sufficiently weakened to permit access to RNase but not enough to drastically reduce cDNA synthesis, consistent with multiple Avd contact points to the RNA. The boundaries of the Avd binding site were probed by scrambling Sp 4-18 and Sp 35-44. The former eliminated RNase protection and severely attenuated cDNA synthesis, while the latter had little effect on either (Figs. S15E-G). Interestingly, with Sp 45-49 scrambled, the RNA was not protected by Avd but was protected by bRT-Avd, and cDNA synthesis was reduced to one-fifth of wild-type (Fig. S15G). This suggests that Avd forms weak contacts to this priming-proximal region, and these contacts are reinforced by bRT. Taken together, these results are consistent with Sp 4-30 forming the Avd binding site, and indicate that Avd, in addition to its role in catalysis, is required for targeting bRT-Avd to the Sp region of DGR RNA.

## DISCUSSION

Adenine-mutagenesis is a hallmark feature of DGRs. We sought to identify the events involved in cDNA synthesis and adenine-mutagenesis through *in vitro* reconstitution of components of the *Bordetella* bacteriophage DGR. An essential step in accomplishing this was isolation of biochemically well-behaved bRT, which required its co-expression with Avd. A second essential step was the purification to homogeneity of *in vitro* transcribed RNA, without which no cDNA synthesis was evident. Once purified, bRT-Avd and DGR RNA recapitulated synthesis of template-primed RNA-cDNA molecules *in vitro*, as observed *in vivo* (19). The correspondence to *in vivo* observations included identical sites of priming (Sp 56) and initiation (*TR* 117) (19). This correspondence further included the level of cDNA synthesis from mutated DGR RNA templates (19). Specifically, substitutions that abrogate or decrease cDNA synthesis *in vivo* did the same *in vitro* (i.e., Sp A56G and Mut4 for the former, and TR U118C and G119A for the latter), and a substitution that has a mild effect *in vivo* also had a mild effect *in vitro* (Sp C55U). Most significantly, these trends in cDNA synthesis closely parallel retrohoming activity (19), providing evidence that the RNA-cDNA molecules observed *in vitro* and *in vivo* are on the path to sequence diversification.

Using the *in vitro* system, we discovered that the bRT-Avd complex was necessary and sufficient for adenine-mutagenesis. bRT by itself had no reverse transcription activity, and required the participation of Avd. Synthesis of adenine-mutagenized cDNA was intrinsic to the bRT-Avd complex, occurring for either DGR or non-DGR RNA templates, and occurring either when primed by the template or an exogenous ODN. Remarkably, misincorporation by bRT-Avd at template adenines occurred with near equal efficiency as proper incorporation. This promiscuity extended to dUTP incorporation, with bRT-Avd displaying an unprecedentedly strong preference for dUTP over TTP (37). In *Bordetella*, the dUTP concentration was measured to be sufficiently low that misincorporation rather than uridine incorporation was the likely pathway for adenine-mutagenesis. The proclivity to incorporate uridines likely reflects the extreme infidelity of bRT-Avd at template adenines, but may also have a role in diversification in organisms or conditions in which the dUTP concentration is substantially higher.

While the RNA template had no apparent role in adenine-mutagenesis, the processivity of bRT-Avd was strictly dependent on the RNA template. This was evident in experiments using exogenous primers. A primer that initiated synthesis from *TR* 117 (P^G117^) resulted in extended cDNA synthesis, but only when the RNA template included intact Sp. When Sp was truncated at Sp 56, only short cDNAs could be primed from *TR* 117 by P^G117^. This was remarkable as P^G117^ annealed to *TR*, quite distant from Sp 56. Likewise, only short cDNAs were synthesized when primed by an ODN from other locations of the DGR RNA or from non-DGR RNA templates. Loss of the *avd* sequence upstream of *TR* had a less dramatic effect than loss of Sp on synthesis primed by P^G117^, but still resulted in a substantial decrease in the synthesis of extended cDNA. These upstream and downstream sequences surrounding *TR* also had a critical impact on template-primed synthesis. Almost the entire Sp region was required for template-primed synthesis, as was 20 nt of *avd.* Taken together, these results suggest that the DGR RNA and bRT-Avd combine to form a ribonucleoprotein (RNP) particle that creates a privileged site for cDNA synthesis from *TR* 117 (Fig. 6E). Recruitment of bRT-Avd by Sp 4-30 along with higher-order structure in Sp, and perhaps *avd*, is likely to position the 2’-OH of Sp 56 as the priming group and *TR* 117 as the templating base at the catalytic site of bRT. Without *avd* and Sp sequences surrounding *TR*, it is likely that no RNP forms or is incorrectly structured such that Sp 56 is unavailable for priming. This would explain why these surrounding sequences are absolutely required for template-primed synthesis of *TR*-cDNA. We further envision that the RNP stabilizes contacts between bRT-Avd and the RNA template, thereby supporting processive polymerization. While an exogenous primer that initiates from *TR* 117 can circumvent the need for template-priming from Sp 56, this would also explain why processive polymerization from such a primer is reliant on *avd* and Sp sequences surrounding *TR*.

Although processive cDNA synthesis can occur from this privileged site either through template- or ODN-primed mechanisms, template-priming provides a clear advantage in obviating the need for additional factors or primer encoded by the DGR (e.g., primase) or the host (e.g., primer). The former may be important for the genomic economy of DGRs, and the latter for the horizontal spread of DGRs to disparate hosts. The existence of a privileged site in the DGR RNA for reverse transcription, beyond providing efficiency for the synthesis of *TR*-cDNA, is also likely to protect the host genome from damage. Genome damage due to retrotransposons, such as LINE and Ty elements, has been documented (42,43). Aberrant reverse transcription of cellular mRNAs by retrotransposon reverse transcriptases results in cDNAs that become incorporated into the genome as pseudogenes via DNA repair and recombination pathways (44). Thus, the functional necessity for the *avd* and Sp regions provides a mechanism for restricting bRT-Avd action to DGR RNA and excluding host RNAs, thereby curtailing the possibility of deleterious pseudogenes arising in host genomes from DGR activity.

Lastly, we found that bRT-Avd is able to carry out RNA cleavage in a manner that is coupled to reverse transcription. However, the functional importance of RNA cleavage remains obscure. We observed cleavage of the core DGR RNA between Sp 57-58, but not the 580 nt DGR RNA, which differs from the core DGR RNA primarily in the sequence length of *avd* upstream of *TR*. RNA cleavage occurs in group II introns as well, and is catalysed in large part by the RNA itself (45). The group II intron maturase protein has a minor role in RNA cleavage, but its reverse transcriptase active site is not required (45), suggesting different mechanisms for RNA cleavage in group II introns and DGRs. Cleavage has also been observed for DGR RNA isolated from *Bordetella*, and here too, cleavage required a catalytically active bRT (19). The implication from this study was that RNA cleavage frees the 3’-OH of Sp 56 to act as the priming site. However, no evidence in this study indicates that RNA cleavage is required for reverse transcription. Our results, on the other hand, provide evidence that RNA cleavage is not required for reverse transcription. By isolating RNA-cDNA molecules and probing for the presence of intact Sp, we found that the priming site at Sp 56 can be internally located within the RNA. We next showed that the 2’-OH of Sp 56, the most chemically plausible site for priming, was required for reverse transcription from both the core and 580 nt DGR RNA templates. This was done by constructing hybrid RNA molecules that had a deoxynucleotide specifically at Sp 56. In the case of the core DGR RNA, we also found that RNA cleavage was dependent on the 2’-OH of Sp 56. Taken together, these observations suggest that reverse transcription is primed by the 2’-OH of Sp 56, and that RNA cleavage between Sp 57-58 follows priming, explaining the dependency of cleavage on the 2’-OH of Sp 56. If RNA cleavage is not required for cDNA synthesis, what is its function? This remains an open question, but our evidence suggests that if RNA cleavage has a functional role, it is likely to be at a step following cDNA synthesis.

In summary, our studies set the stage for addressing the next steps in mutagenic retrohoming as well as elucidating the structural basis for the uniquely selective promiscuity of bRT-Avd.

## SUPPLEMENTARY DATA

Tables S1-S3

Figs. S1-S15

Supplementary Data are available at NAR online.

## ACKNOWLEDGEMENT

We thank S. Naorem and H. Guo for sharing results prior to publication. We also thank A. Robart, N. Toor, S. Joseph, and G. Ghosh for discussion and helpful suggestions.

